# Decreased electrocortical temporal complexity distinguishes sleep from wakefulness

**DOI:** 10.1101/691006

**Authors:** Joaquín González, Matias Cavelli, Alejandra Mondino, Claudia Pascovich, Santiago Castro-Zaballa, Pablo Torterolo, Nicolás Rubido

## Abstract

In most mammals, the sleep-wake cycle is constituted by three behavioral states: wakefulness (W), non-NREM (NREM) sleep, and REM sleep. These states are associated with drastic changes in cognitive capacities, mostly determined by the function of the thalamo-cortical system. The intra-cranial electroencephalogram or electocorticogram (ECoG), is an important tool for measuring the changes in the thalamo-cortical activity during W and sleep. In the present study we analyzed broad-band ECoG recordings of the rat by means of a time-series complexity measure that is easy to implement and robust to noise: the Permutation Entropy (PeEn). We found that PeEn is maximal during W and decreases during sleep. These results bring to light the different thalamo-cortical dynamics emerging during sleep-wake states, which are associated with the well-known spectral changes that occur when passing from W to sleep. Moreover, the PeEn analysis allows to determine behavioral states independently of the electrodes’ cortical location, which points to an underlying global pattern in the signal that differs among the cycle states that is missed by classical methods. Consequently, our data suggest that PeEn analysis of a single EEG channel could allow for cheap, easy, and efficient sleep monitoring.

## Introduction

The sleep-wake cycle is a critical physiological process and one of the most preserved biological rhythms through evolution.^1^ This cycle is composed of different states, commonly distinguished by their electro-physiological signatures and behavioral characteristics. These states correspond to wakefulness (W), non-rapid eye movement (NREM) sleep, and rapid eye movement (REM) sleep. W and sleep are associated to different brain functional states, which can be captured by electroencephalographic (EEG) signals containing a broad frequency spectrum. Accompanying the electrocortical differences among the states, the cognitive capacities drastically change during the cycle. Fundamentally, consciousness is lost during deep NREM sleep, emerging in an altered fashion during REM sleep. Altered states of consciousness can also arise during special normal states, such as during lucid-dreams,^2^ or under toxic or pathological conditions, such as the states induced by psychedelic drugs or psychosis.^3, 4^

Cognitive states are mostly determined by the function of the thalamo-cortical system.^1^ Part of this neuronal processing can be accurately measured by intra-cranial elecroencephalogram (EEG), known as electrocorticogram (ECoG). Due to the complex nature of the standard EEG and ECoG signals, traditional methods employed in neuroscience have divided the complex spectrum of the signal into frequency bands,^3, 5–9^ and analyzed its changes during different cognitive functions,^4, 7^ and sleep states.^5, 6, 8^ These methods only describe particular characteristics of the recorded signals and do not account for the complex nature of the cortical electric potentials. In contrast, the field of non-linear dynamics has developed measures and models that account for the complexity of the systems and their emerging interactions.^10–13^ These properties are fundamental for the characterization of the thalamo-cortical function and for the emergence of consciousness.^14^

A general approach to study time-signals is the characterization of their randomness; for example, by means of the Shannon entropy (SE),^15^ which measures the average unpredictability of a signal. However, SE requires a random source and an invariant probability distribution (which is typically unknown), also is affected by noise, by measurement precision, and by data length. All these elements are relevant when dealing with real-world signals. In order to find a similar non-linear measure to quantify unpredictability from real-world data, Bandt and Pompe^10^ introduced the Ordinal Pattern (OP) analysis, allowing to encode any signal into OPs and approximate its SE. This approximation is known as Permutation Entropy (PeEn). In contrast to other methods, PeEn is a time-series complexity measure that is simple to implement, is robust to noise and short time-series, and works for arbitrary data sets.^13, 16, 17, 20–26^ In particular, it has been shown that PeEn applied to EEG signals captures different states associated with the level of consciousness, both during anesthesia^26–29^ and sleep.^30, 31^ Hence, in order to study the thalamo-cortical function during W and sleep, PeEn is a practical and reliable method, where results can be understood from primary principles, and can be related to the signal characteristics.

Previous works have analyzed PeEn in standard EEG recordings.^27–31^ However, EEG signals have frequency limitations due to scalp-filtering, are often recorded with low sampling rates, and pre-acquisition filters are commonly applied. In addition, as the recording electrodes are placed above the scalp’s skin, other sources can interfere with the cortical neural activity (e.g., muscular activity).^32^ These limitations exclude the possibility of considering high-frequency oscillations; for example, *γ* frequency band (30−100 *Hz*), which is known to vary substantially during the sleep-wake cycle and is an active field of research in Neuroscience.^3, 5–9, 33, 34^ Thus, PeEn analysis of standard EEG signals overlooks the significance of the broad frequency spectrum in relation with the thalamo-cortical function, its cognitive counterpart, and its different electrocortical oscillations. Consequently, it is still uncertain whether previous results hold when considering ECoG measurements and whether these results would depend on cortical location, frequency content, or PeEn parameters.

In the present study, we characterized the PeEn of recordings from freely moving rats during W and sleep. We found that ECoG’s PeEn is maximal during W and decreases during both sleep states. Moreover, we noted that these results are independent of the cortical location (namely, the electrodes placement), pointing towards a global cortical pattern for each sleep-wake state that is captured by the PeEn analysis but is missed by classical methods.

## Results

### Permutation Entropy during Wakefulness and Sleep

Figure 1a shows examples of polysomnographic recordings obtained from electrodes placed directly above the cortex of a representative rat. Electrode locations are shown on the left panel and the intra-cranial polysomnographic recordings for W (blue), NREM (green), and REM sleep (red) states are shown on the right panels, which have been distinguished by means of the standard sleep scoring criteria (see Methods). In Fig. 1b we show, for the same animal, the hypnogram (top), as well as the spectrogram (middle) and PeEn values (bottom) processed from the ECoG recorded with the V2r electrode (with a sampling rate of 1024 *Hz* and *D* = 3 embedding dimension for the PeEn analysis; see Methods for details). The hypnogram shows the standard sleep scoring, the spectrogram shows the ECoG frequency content, and the PeEn quantifies its complexity; namely, its randomness or unpredictability. Maximal PeEn values were achieved during W, PeEN values decreased during NREM sleep and reached minimum values during REM sleep. This result states that the ECoG becomes more predictable – less random – during sleep, especially during REM sleep. More importantly, we found that PeEn is able to detect transitions between behavioral states, which are typically difficult to be noticed from raw data or from the spectrogram. For example, the power spectrum in Fig. 1b (middle panel) shows similarities between W and REM sleep, but PeEn values are drastically different between these states (W epoch at 11 to 13 minutes and REM sleep epoch at 22 to 24 minutes). The average PeEn values for all rats, behavioral states, and cortical locations are shown in Fig. 1c and table. 1, there are significant differences among behavioral states for all recorded neocortical regions; in the Olfactory Bulb (OBr, archicortex) there was a tendency to decrease (*p* = 0.056) when REM was compared to NREM sleep (first row in Table 1). Consequently, the PeEn of the ECoG characterizes and follows the state transitions regardless of the electrode’s location.

**Figure 1.**
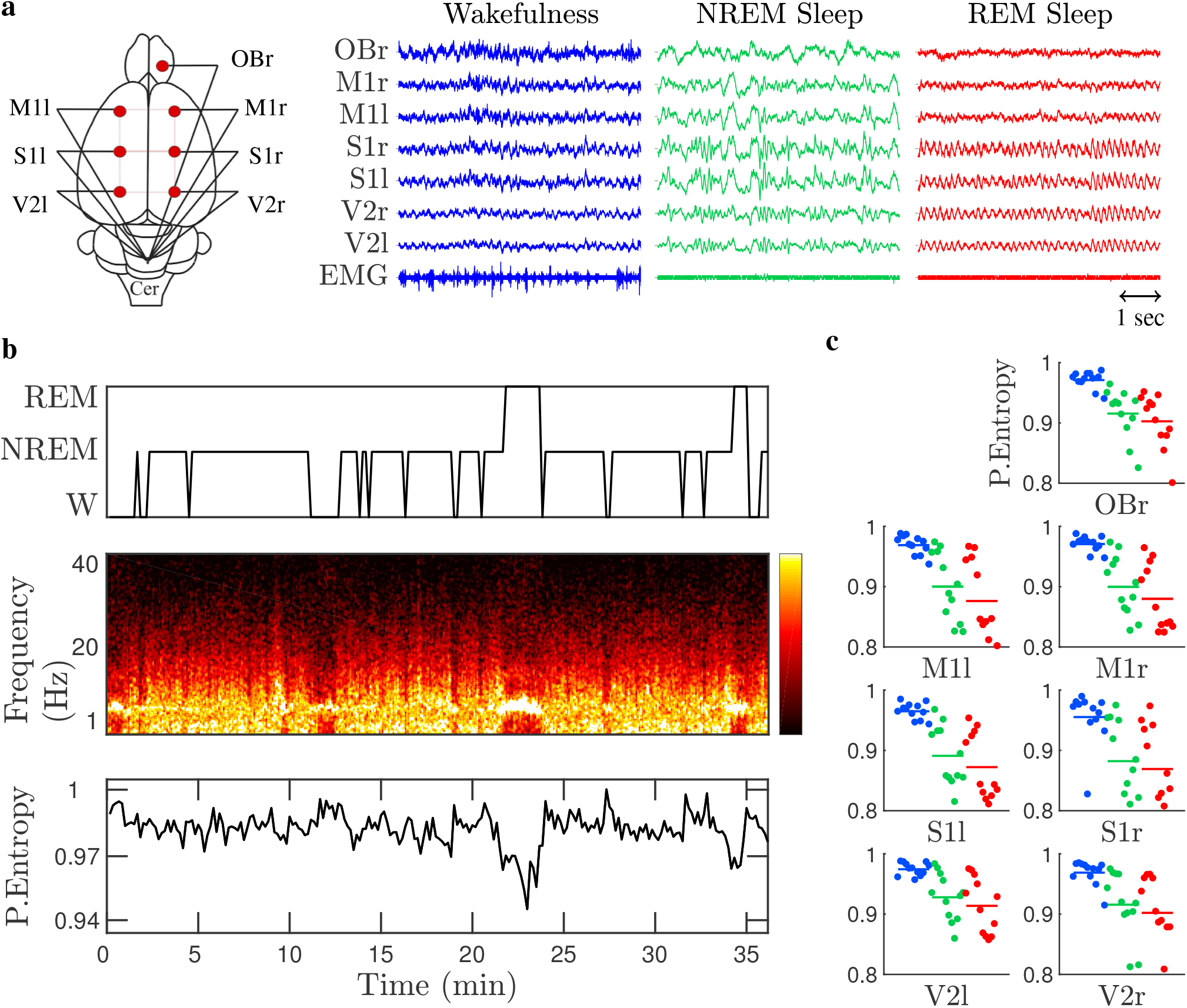
Permutation Entropy (PeEn) during wakefulness and sleep. Panel **a** shows a schematic representation of the placement of the 7 electrodes across the cortex, representative ECoG (5 second window referenced to the cerebellum and the neck electromyogram (EMG), for each sleep-wake state: wakefulness (blue), NREM (green) and REM sleep (red). From top to bottom, Olfactory Bulb (OBr), right and left Primary Motor (M1r/M1l), Primary Somatosensory (S1r/S1l), and Secondary Visual (V2r/V2l) cortices. Using 30 *s* sliding windows for the V2r electrode, panel **b** shows the hypnogram (top) with the visually scored sleep states, the power spectral density (middle) with yellow indicating high power, and PeEn analysis (bottom) for embedding dimension *D* = 3. Panel **c** gathers the time-averaged PeEn values (for embedding dimension *D* = 3) for 12 rats, differentiating each cortex electrode and sleep state (colour code as in panel **a**). Namely, each dot in panel **c** corresponds to the time-averaged PeEn value of each rat and cortex, where the horizontal bars are the population mean (the statistic is shown in Table 1).

**Table 1.** Statistical comparisons between PeEn values during sleep and wakefulness. Each row corresponds to a different cortical location, as shown in Fig. 1a. Data was evaluated by repeated measures ANOVA and Bonferroni *post-hoc* test. These results correspond to encoding the electro-corticographic signals with *D* = 3 and 1024 *Hz* sampling frequency (see Methods for details).

| Electrode | F | DF | W-NREM<br>(pValue) | W-REM<br>(pValue) | NREM-REM<br>(pValue) |
| --- | --- | --- | --- | --- | --- |
| OBr | 40.2 | 2,11 | 0.0003 | 0.0002 | 0.056 |
| M1r | 37.68 | 2,11 | 0.0005 | 0.0002 | 0.028 |
| M1l | 29 | 2,11 | 0.0014 | 0.0007 | 0.007 |
| S1r | 32.1 | 2,11 | 0.0011 | 0.0007 | 0.046 |
| S1l | 41.18 | 2,11 | 0.0004 | 0.0002 | 0.005 |
| V2r | 23.09 | 2,11 | 0.0043 | 0.0017 | 0.009 |
| V2l | 24.92 | 2,11 | 0.0022 | 0.0017 | 0.036 |

### Permutation entropy dependence on the embedding dimension

We characterized how PeEn reflects the ECoG temporal complexity during W and sleep. In order to do this characterization, we modified the embedding dimension *D*, which is the parameter that sets the ordinal pattern (OP) length encoding the ECoG signal. Specifically, each OP captures the relationship between the relative amplitudes (ranking its values) inside a *D*-sized non-overlapping time-window of the signal (see Methods for details on the encoding procedure). Hence, changing *D* modifies the time scale of the ECoG being analyzed and the resultant PeEn calculation. The larger the OP, the more details are obtained from the signal; thus, the less random the signal becomes and the smaller its PeEn value. For example, Fig. 2 shows that the PeEn values consistently decrease as we increased the embedding dimension from *D* = 2 to *D* = 4, either for W or sleep. Overall, averaged PeEn values during W were larger than during sleep for all *D* s. Also, the fact that PeEn variability is minimal for *D* = 2 is because this dimension has the greatest sensitivity to random fluctuations: i.e., it mainly captures noise.

**Figure 2.**
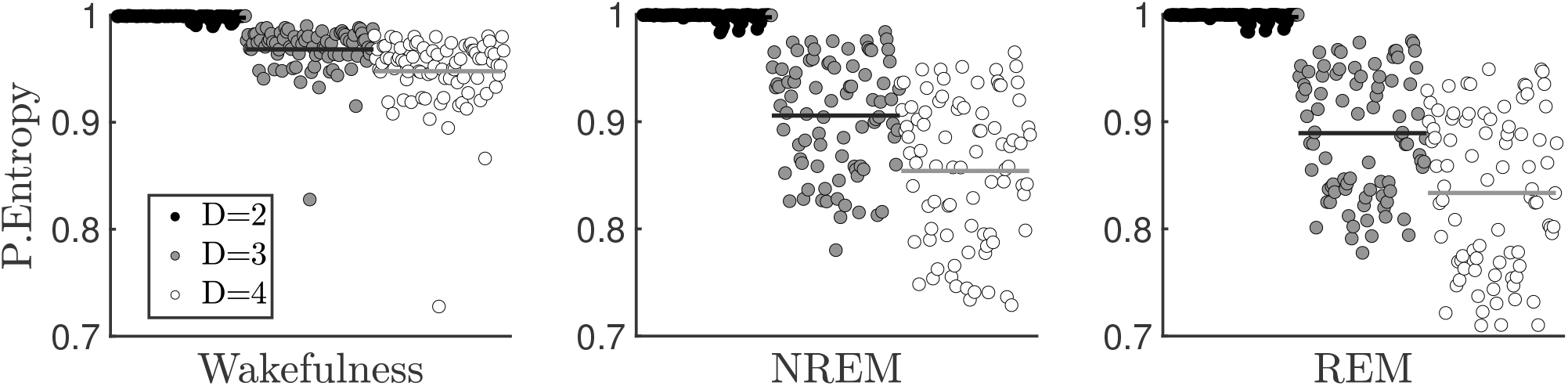
Permutation Entropy (PeEn) of electro-corticograms (ECoG) as a function of the embedding dimension. The PeEn values are normalized according to the maximum possible entropy for each embedding dimension, *D*; namely, by log(*D*!). From left to right, wakefulness (W), non-rapid eye movement (NREM) and rapid eye movement (REM) sleep are shown. Symbols represent each PE value for all electrode locations (*n* = 7) and animals (*n* = 12) [as shown in Fig. 1c], when using *D* = 2 (black), *D* = 3 (grey), or *D* = 4 (white) for the ordinal pattern encoding of the ECoG signals recorded at a sampling frequency of 1024 *Hz*. The horizontal lines represent the population and electrode location average for the respective embedding dimensions.

### Permutation entropy relationship with the frequency spectrum

Standard EEG and ECoG analyses often involve decomposing the signals into frequency bands, which are defined considering different physiological processes. In order to associate the PeEn values with the different frequency bands, we performed successive down-samples to the ECoG signals. This process allows for the ordinal patterns to capture lower frequency components of the ECoG signal, while keeping constant the embedding dimension, *D*.

We down-sampled the ECoG from a sampling rate of 1024 *Hz*, halving it down to 64 *Hz*; thus, changing the maximum frequency resolution from 512 *Hz* to 32 *Hz*. By doing this, the PeEn value changed, revealing its relationship to the frequency spectrum. Fig. 3a shows averaged PeEn using *D* = 3 for all rats and electrode locations, for W (blue), NREM (green) and REM sleep (red). The shaded areas indicate the mean ± standard error of these averages, showing that the states differentiate in average significantly (the exact statistics are exhibited in Table. S.1 at the Supplementary Material). Average PeEn values for the ECoGs during REM sleep are larger than NREM sleep until the maximum frequency resolution increases beyond 128 *Hz*, remaining lower than W values for all frequencies. Although Fig. 3a shows only the PeEn values for *D* = 3, we obtained similar results for larger embedding dimensions (data not shown). The relationship between the PeEn values and the sampling frequency can be further understood by comparing these results with the power spectral density (PSD) analysis shown in Fig. 3b. In general, the PSD of a signal is the probability distribution function of its frequency content; namely, the degree of presence that each frequency component has in the signal. As we down-sampled the ECoG signals, as in Fig. 3a, the higher frequencies are cut-off from the PSD [Fig. 3b]. For the higher frequencies, i.e., > 200 *Hz*, there is a large difference in the PSD value between W and sleep. However, when lower frequencies are considered, REMs PSD increased above NREMs. Note that below 200 *Hz* REMs lower frequencies become more relevant (Fig. 3b) and PeEn (Fig. 3a) values are larger than NREM sleep and closer to W.

**Figure 3.**
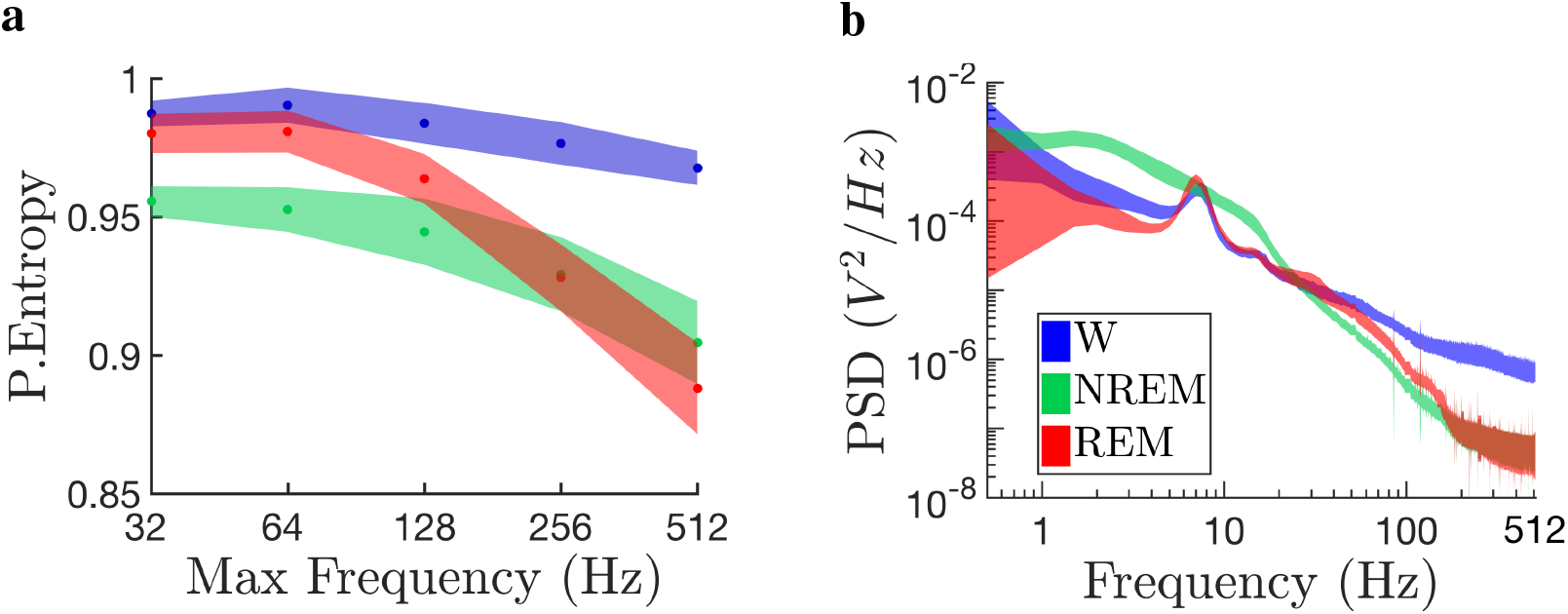
Permutation entropy (PeEn) relationship with frequency content and power spectrum (PSD). Panel **a** [Panel **b**] shows the averaged PeEn [averaged PSD], values as a function of the electro-corticograms (ECoG) maximum frequency resolution [frequency components] for each sleep-wake state (colour codes are as indicated in panel **b**’s inset). Both averaged quantities are the resultant mean value of the 7 cortical electrode placements for all 12 rats analyzed (shown in Fig. 1). Shaded areas depict the standard error of the mean in Panel **a** and twice the standard error of the mean in Panel **b**. The maximum frequency resolution is the ECoGs sampling frequency divided by 2, according to the Nyquist-Shannon criterion.

### Ordinal Pattern Probability Distributions

In addition to the Entropy quantification, we considered the qualitative differences and variations appearing in the OP probability distributions during W and sleep. The OP distributions shown in Fig. 4 quantify the relative frequency of appearance that each OP has in the encoded ECoG signal; namely, the OP probability. Figure 4a shows the 6 possible OPs when the embedding dimension is *D* = 3 (top panel), and the resultant OP probability distribution we found from the ECoGs in each sleep-wake state (bottom panel). Similarly, Fig. 4b shows the OPs and OP distribution for *D* = 4. It is readily observed from both panels that the increasing or decreasing OPs (i.e., labels 1 and 6 in Fig. 4a and labels 1 and 24 in Fig. 4b) have a larger probability of occurrence, irrespective of the sleep-wake state or the embedding dimension (results hold for larger *D* – not shown). We note that, in spite of having qualitatively similar distributions for *D* = 4, other OPs start to emerge, such as labels 7 and 18, which are modified versions of labels 1 and 24, respectively. Nevertheless, these OPs are not statistically significant, since they fall within the null hypothesis confidence interval (signaled by the shaded grey areas in both panels). On the contrary, the OP distribution in Fig. 4a for *D* = 3 during sleep (red and green) significantly departs from the null hypothesis, which corresponds to the uniform distribution (shaded grey area).

**Figure 4.**
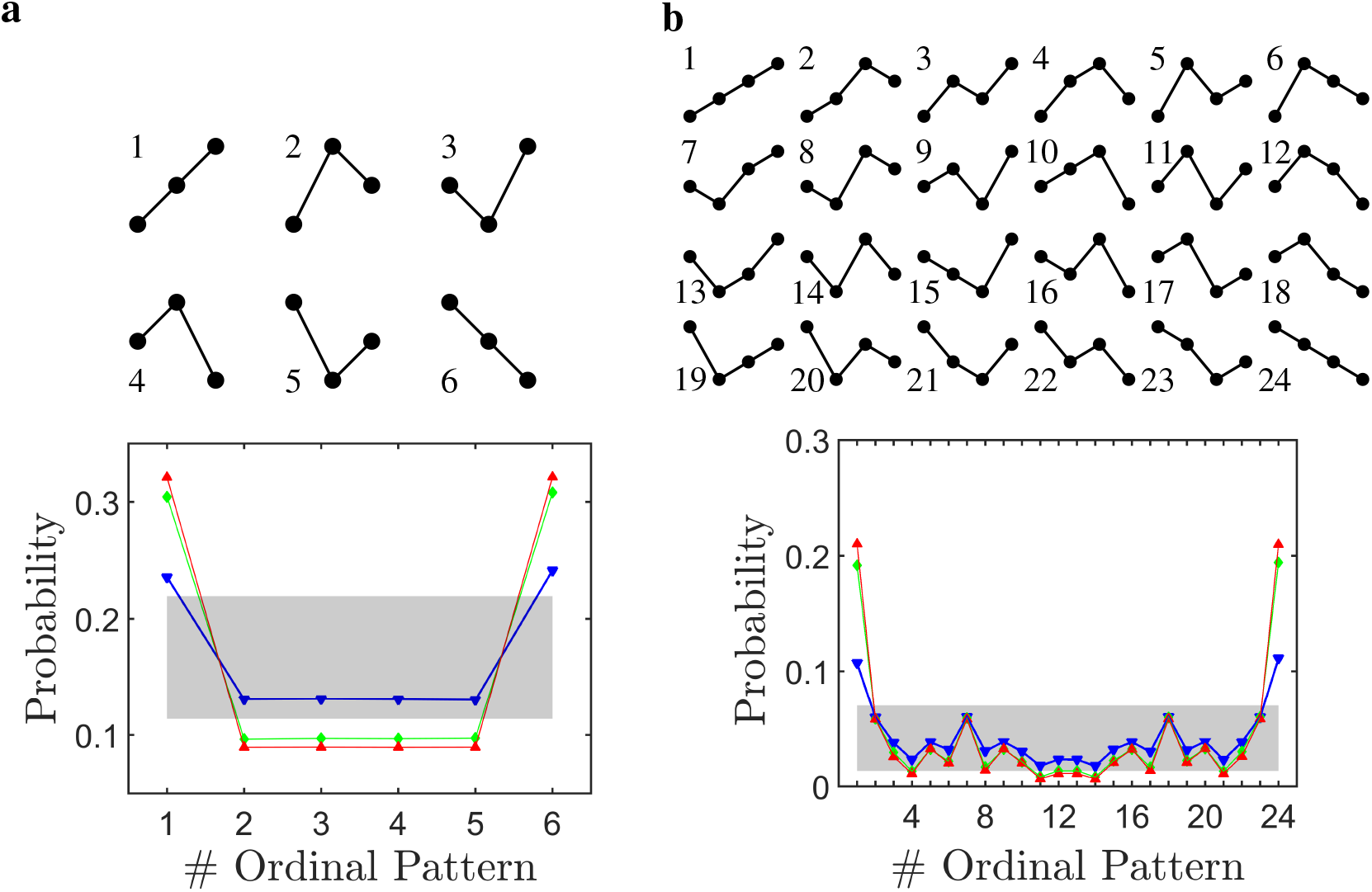
Ordinal Pattern (OP) probability distributions during wakefulness and sleep. The OP probability distributions shown in the bottom panels correspond to the rat population and electrode location average distributions for each sleep-wake state: Wakefulness (blue), NREM (green) and REM sleep (red). The grey areas show the null hypothesis region with a 95, 4% confidence, which correspond to the uniform OP distribution with twice the standard error of the mean (i.e., *p*_*NH*_ ± 2*σ_NH_*). Panel **a**[Panel **b**] shows the possible OPs for embedding dimension *D* = 3 [*D* = 4].

**Figure 5.**
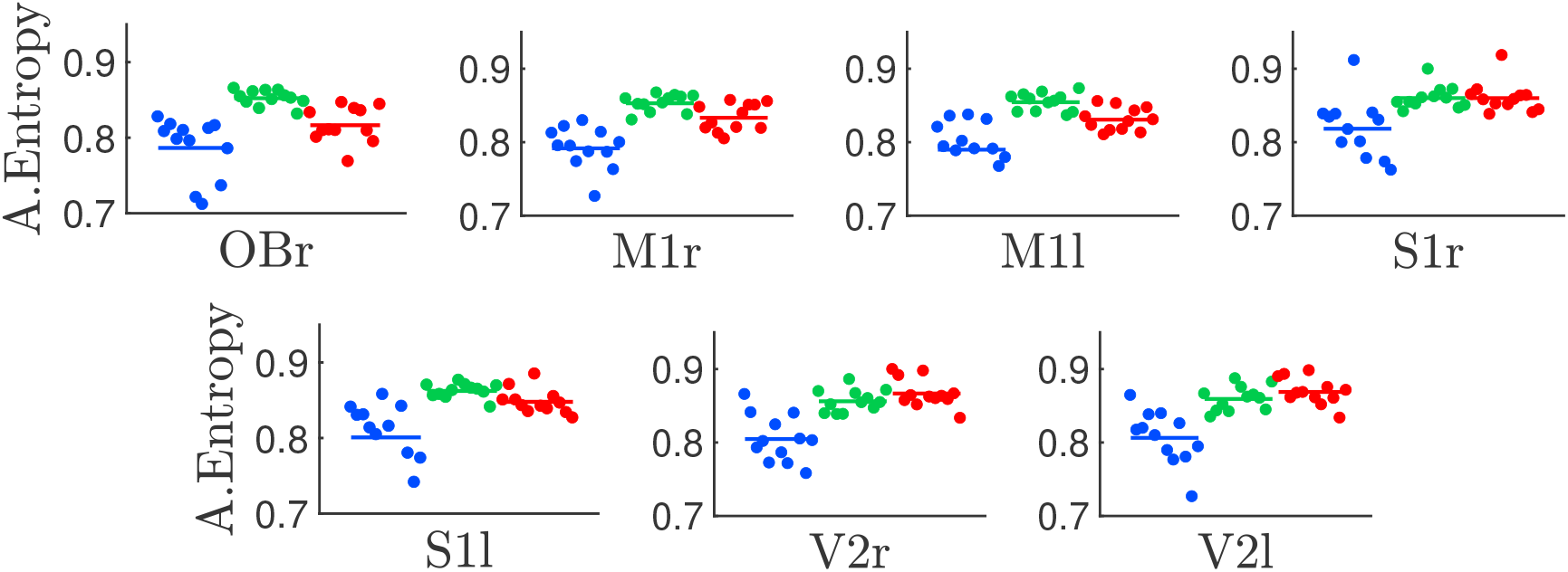
Amplitude entropy during wakefulness and sleep. From left to right and top to bottom, the panels show the entropy values calculated from the electro-corticographic (ECoG) histograms coming from the different cortical locations shown in Fig. 1; i.e., olfactory bulb, right and left motorsensory, somatosensory, and visual cortices, respectively. The colour code signals the sleep-wake cycle states (W, blue; NREM, green and REM sleep, red) and the symbols and horizontal lines represent the same as in Fig. 1c. The entropy values were calculated from the ECoG amplitude histograms using 18 bins. The sampling rate was 1024 *Hz*. The statistic is shown in the Supplementary Material (see Table. S.2).

### Comparison with classical amplitude encoding

Classical analysis of time-series uses the probability distribution function of the signal; namely, the signal is encoded using a histogram of its amplitudes. This process discards the information coming from the signal’s time stamps; in other words, the amplitude time-dependence. We compared the entropy values using histograms with 18 bins of the ECoG, where the results are shown in Fig. 1c. As can be directly observed, there were some differences among the sleep states (either NREM or REM), but there were no consistent global pattern and no single electrode was able to differentiate between all sleep-wake states (see Table. S.2). Moreover, these results remain practically invariant when using larger number of bins (data not shown).

## Discussion

In this work, we described that the collective cortical activity measured by ECoG in male adult rats fluctuates between periods of high temporal complexity during W, and periods of low temporal complexity during sleep (see Fig. 1b and **c**). These ECoG complexity variations reflect the differences in the thalamo-cortical function between sleep-wake states. Consequently, our results strongly support and extend studies in human that carried out PeEn analyses and other complexity measures in standard EEG recordings.^27–31, 35^

We also showed that PeEn profile during W and sleep did not change according to the cortical recording site, reflecting a common micro-structure motif and a dynamical behavior which are independent from the origin of the cortical signal. The average randomness of these micro-structure patterns distinguished sleep from W, regardless of the changes in the embedding dimensions employed in PeEn analysis (Fig. 2) and the sampling frequencies considered (Fig. 3a). These results suggest the use of PeEn as a quantitative tool for understanding thalamo-cortical dynamics during various physiological conditions, the influence of psychoactive drugs, or pathological conditions.

A strong benefit from PeEn analyses is that PeEn variations across states can be explained by dynamical systems theory;^10, 16, 18^ conversely, with other techniques the tractability is lost (such as, machine-learning approaches). During NREM sleep, the neuro-modulation coming from the activating systems drastically decreases, which favors the occurrence of slow *δ* waves (1 − 4 *Hz*) and sleep spindles (9 − 12 *Hz*) in the thalamus and cortex.^1^ This means that, as the cortex transitions from W to sleep, the higher-frequency cortical patterns (complex signals) decrease, while lower-frequency oscillations (less complex signals) rise (Fig. 3b). As a consequence, the OPs probability distribution becomes less uniform (in other words more predictable). Specifically, we found that strictly increasing or decreasing OP motifs are strongly favored in all ECoG signals, particularly during sleep (Fig. 4), making the remaining OP motifs to appear less frequently.

Surprisingly, we found that NREM’s PeEn is larger than REM’s PeEn, which contradicts previously reported results.^30, 31^ owever, we showed that as the frequency content of the ECoG varies, the PeEn changes its value. For example, Fig. 3a exhibits that as we down-sampled the ECoG, REM sleep PeEn becomes larger than NREM’s PeEn; these low sampling rates are similar to those used in previous studies.^30, 31^ Moreover, we analyzed the results from the PeEn as a function of the maximum frequency resolution (Fig. 3a) in conjunction with those from the power spectral density (PSD) of the ECoG (Fig. 3b). We observed that the rise in REM’s PeEn as the frequency content decreased follows the variations in the PSD. Specifically, REM sleep presents larger power than NREM sleep around and below 120 *Hz* corresponding to the high frequency oscillations and gamma band oscillations.^5, 6^ This means that when higher frequencies are cut-off, the PSD slope significantly increases approaching a more uniform frequency distribution, which corresponds to a more complex time-series. These analyses reveal that PeEn results depends on the frequency content of the signal. In particular, REMs temporal complexity resembles W when gamma oscillations are captured by the PeEn.

One of the main differences between W and sleep, is that muscle tone and movements are mainly absent during sleep (specially during REM sleep). In this regard, the power of the higher frequencies of the spectrum is significantly higher during W than during sleep. It is possible that exists a contribution of muscular activity on the ECoG (by volume conduction) on the high frequency bands (> 100 *Hz*), as supported by experimental evidence.^32^ Hence, the high values of PeEn during W could be determined by the muscle electrical activity (produced mainly by the muscle tone, because epochs with movement artifacts were discarded from the analysis) that inevitably pollutes the ECoG. Nevertheless, the PeEn values remained constant during W following downsampling, in spite of the fact that higher frequencies were bypassed and the contribution from muscle tone became less relevant. Still, more research is needed in order to quantify the weight of the muscle artifact in the PeEn results.

As a final remark, in clinical settings, sleep classification is usually performed by visual or spectral analysis of the EEG’s waveform at different recordings sites, together with the examination of the EMG and electroculogram. In contrast, our data suggest that robust and reliable automatic sleep monitoring could be achieved by means of PeEn analysis. Hence, PeEn analysis of a single EEG channel may provide a cheap and efficient procedure for sleep monitoring.

## Methods

### Experimental Animals

All experimental procedures were conducted in agreement with the National Animal Care Law (No. 18611) and with the “Guide to the care and use of laboratory animals” (8th edition, National Academy Press, Washington DC, 2010). Furthermore, the Institutional Animal Care Committee approved the experiments, where 12 Wistar adult rats were maintained on a 12 − *h* light/dark cycle (lights on at 07 : 00*h*) with food and water freely available. The animals were determined to be in good health by veterinarians of the institution. We took adequate measures to minimise pain, discomfort, and stress in the animals, and all efforts were made to use the minimal number of animals necessary to obtain reliable scientific data.

### Surgical Procedures

The animals were chronically implanted with electrodes to monitor the states of sleep and W. We employed similar surgical procedures as in previous studies.^5, 6^ Anaesthesia was induced with a mixture of ketamine-xylazine (90 *mg/kg*; 5 *mg/kg* i.p., respectively). The rat was positioned in a stereotaxic frame and the skull was exposed. To record the ECoG, stainless steel screw electrodes were placed on the skull above motor, somatosensory, visual cortices (bilateral), the right olfactory bulb, and cerebellum, which was the reference electrode (see Fig. 1a and Table. 2). In order to record the EMG, two electrodes were inserted into the neck muscle. The electrodes were soldered into a 12-pin socket and fixed onto the skull with acrylic cement. At the end of the surgical procedures, an analgesic (Ketoprofen, 1 *mg/kg*, s.c.) was administered. After the animals had recovered from these surgical procedures, they were left to adapt in the recording chamber for 1 week.

**Table 2.** Electrode Location. Schematic representation and electrode locations. All coordinates are referenced to *Bregma* (Lateral: 0, Antero-posterior: 0) according to Paxinos and Watson 2006^36^

| Electrode | OB<sub>r</sub> | M1<sub>r</sub> | M1<sub>l</sub> | S1<sub>r</sub> | S1<sub>l</sub> | V2<sub>r</sub> | V2<sub>l</sub> |
| --- | --- | --- | --- | --- | --- | --- | --- |
| Antero-Posterior | +7.5mm | +2.5mm | +2.5mm | -2.5mm | -2.5mm | -7.5mm | -7.5mm |
| Lateral | +1.25mm | +2.5mm | -2.5mm | +2.5mm | -2.5mm | +2.5mm | -2.5mm |

### Experimental Sessions and Sleep Scoring

Animals were housed individually in transparent cages (40 × 30 × 20 *cm*) containing wood shaving material in a temperature-controlled (21 — 24^◦^ C) room, with water and food *ad libitum*. Experimental sessions were conducted during the light period, between 10 AM and 4 PM in a sound-attenuated chamber with Faraday shield. The recordings were performed through a rotating connector, to allow the rats to move freely within the recording box. Polysomnographic data were amplified (*X* 1000), acquired and stored in a computer using Dasy Lab Software employing 1024 *Hz* as a sampling frequency and a 16 bits AD converter. The states of sleep and W were determined in 10 s epochs. W was defined as low voltage fast waves in the motor cortex, a strong theta rhythm (4 — 7 *Hz*) in the visual cortices, and relatively high EMG activity. NREM sleep was determined by the presence of high voltage slow cortical waves together with sleep spindles in motor, somatosensory, and visual cortices associated with a reduced EMG amplitude; while REM sleep as low voltage fast frontal waves, a regular theta rhythm in the visual cortex, and a silent EMG except for occasional twitches. An aditional second visual scoring was performed in 30 s epochs where artifacts and transitional states were discarded.

### Ordinal Pattern encoding

In order to quantify the EEGs’ randomness, we encoded the time-series into ordinal patterns (OPs) following Bandt and Pompe method.^10^ The encoding involves dividing a time-series, {*x*(*t*)*, t* = 1*, …, T*}, into ⌊(*T — D*)*/D*⌋ non-overlapping vectors, where ⌊*y*⌋ denotes the largest integer less than or equal to *y* and *D* is the vector’s length, which is much shorter than the time-series length (*D ≪ T*). Then, each vector is classified according to the relative magnitude of its *D* elements. The classification was done by determining how many permutations are needed to order its elements increasingly; namely, an OP is associated to represent the vector’s permutations. For example, for *D* = 2, the time-series would be divided into vectors containing two consecutive values, such as {*x*(*t*_*i*_)*, x*(*t*_*i*+1_)}, that are non-overlapping (the next vector to {*x*(*t*_*i*_)*, x*(*t*_*i*+1_)} is the {*x*(*t*_*i*+2_)*, x*(*t*_*i*+3_)} vector, where *t*_*i*_ is the *i*-th time stamp). These vectors have only two possible OPs for any time *t*_*i*_: either *x*(*t*_*i*_) < *x*(*t*_*i*+1_) or *x*(*t*_*i*_) *> x*(*t*_*i*+1_), which correspond to making 0 permutation or 1 permutation, respectively. It is worth noting that the number of possible permutations increases factorially with increasing vector length; i.e, for vectors of length *D* there are *D*! possible OPs. In particular, we labeled the OPs as the number of permutations plus one; hence, our OPs are labeled by means of integers, *α*, that range from *α* = 1 to *α* = *D*!. For *D* = 2, *α* = 1 or 2. Similarly, for *D* = 3, the OPs *α* = 1 and *α* = 2 correspond to having a vector from the time-series with 3 values ordered as *x*(*t*_*i*_) < *x*(*t*_*i*+1_) < *x*(*t*_*i*+2_) and *x*(*t*_*i*_) < *x*(*t*_*i*+2_) < *x*(*t*_*i*+1_), respectively, but there are 4 more possibilities (for *D* = 3, *α* = 1, …, 3! = 1, …, 6). Small noise fluctuations were always introduced into the time-series in order to remove degeneracies; i.e., avoid the cases where, for example, *x*(*t*_*i*_) = *x*(*t*_*i*+1_).

### Randomness quantification

Shannon entropy (SE) is a quantity used in Information theory to quantify the average randomness (information content) of a signal. It is defined as^15^ *H*(*S*) = −∑_*α ∈ S*_ *p*(*α*) log[*p*(*α*)], where *p*(*α*) is the probability of finding symbol *α* in the signal (among the set of symbols *S*) and the summation is carried over all possible symbols. In other words, SE shows that *H*(*S*) is the average value of log(1*/p*) with respect to an alphabet *S*. Hence, in order to find *H* for any real-valued time-series, we need to transform the time-series into a symbolic sequence. When using OPs, the resultant symbolic sequence has a finite number of symbols; i.e., the alphabet, which is given by the OP’s length *D* and holds *D*! = # {*S*} possible symbols. For bin histograms, the number of possible symbols depends on the number of bins, *N*_*b*_, used to create the time-series histogram, which is another way of encoding any bounded time-series into a finite set of values. In order to compare entropy values coming from OPs or bins, we need to set both quantities such that the probabilities involved in the summation of *H*(*S*) are found with identical statistics. For example, when using non-overlapping OPs with *D* = 3, there are *D*! = 6 possibly different symbols in an encoded time-series of length *T*, which accounts to ∼*T/D* total encoded symbols.

We highlight that the number of bins we chose corresponds to making an amplitude encoding that has the same statistical average as the OP encoding with dimension, *D*. Namely, a signal with *T* time-stamps, is encoded by non-overlapping OPs into a symbolic sequence of length *S* = ⌊(*T* − *D*)*/D*⌋ ≃ *T/D*, where ⌊·⌋ indicates the smaller integer closer to the argument. The resultant range for the symbolic sequence distribution is *D*!, which is the different OP possibilities. This means that a length *T* time series has an OP statistical average of *S/D*! ≃ *T/D* × *D*!. On the other hand, the statistical average for histograms with *N*_*b*_ bins of the same time-series is *T/N_b_*. Consequently, in order to have the same statistical average per bin and be able to compare the results, we need to set *N*_*b*_ = *D* × *D*!, which for *D* = 3 corresponds to having *N*_*b*_ = 3 × 6 = 18 bins.

### Power Spectral Density and Statistical Analysis

The power spectral densities were performed using the *pwelch* function on MATLAB by employing the following parameters: window = 30*s,* noverlap = [], fs = 1024, nfft = 1024. These parameters correspond to 30 second sliding windows with half windows overlap, a *f*_*s*_ = 1024 *Hz* sampling frequency and a frequency resolution of 1*Hz*. On the other hand, the statistics for each ECoG PeEn calculations were based on non-overlapping windows of size *D* × *D*! × *N*_*S*_, where *N*_*S*_ = 200 is the statistical average we use for our null-hypothesis. Namely, our null-hypothesis is a Bernoulli process where each ordinal pattern of size *D* has an equal probability of appearance, *p*_*NH*_ = 1*/D*!, and a standard error of the mean 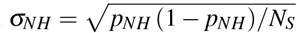. For example, for ordinal patterns with *D* = 3, the non-overlapping windows contained 3 × 6 × 200 = 3600 data points, which accounts to approximately 3.5 seconds at a *f*_*s*_ = 1024 *Hz* sampling frequency. The null-hypothesis in this case has an average probability *p*_*NH*_ = 1*/*3! = 1*/*6 ≃ 0.167 and a standard error of the mean 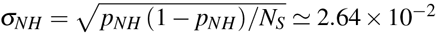, which makes its confidence interval *p*_*NH*_ ± 2*σ_NH_* narrow and the statistical significance of the PE results robust. For the state comparisons, we verified that PeEn distributes normally through Lilliefors test, and then applied a repeated measures ANOVA together with the Bonferroni *post-hoc* test and *p* < 0.05 in order for the result to be considered significant.

## Author Contributions

J.G, N.R and P.T designed the study. J.G, A.M and M.C performed the experiments and data collection. J.G and M.C carried out the data analysis and N.R developed the software. J.G, M.C, A.M, S.C, C.P, P.T and N.R were involved in the discussion and interpretation of the results. J.G, N.R and P.T wrote the manuscript. P.T and N.R provided the financial support. All the authors participated in the critical revision of the manuscript, added important intellectual content, and approved the definitive version.

## Acknowledgements

This study was supported by the “Programa de Desarrollo de Ciencias Básicas”, PEDECIBA; Agencia Nacional de investigación e innovación (ANII), (FCE 1 2017 1 136550) and the “Comisión Sectorial de Investigación Científica” (CSIC) I+D-2016-589 grant from Uruguay. N.R. acknowledges the CSIC group grant “CSIC2018 - FID 13 - Grupo ID 722”.

## Additional Information

**Competing financial interests:** The authors declare no competing financial interests.

## Supplementary Material

**Table S.1.** PeEn and frequency content statistical summary. This table provides the statistics for Fig. 3a.

| Max Frequency | F | DF | State Comparison | pValue |
| --- | --- | --- | --- | --- |
| 512 Hz | 225.58 | 2, 166 | W-NREM | $9.5605e-10$ |
| | | | W-REM | $9.5605e-10$ |
| | | | NREM-REM | $9.5605e-10$ |
| 256 Hz | 239.23 | 2, 166 | W-NREM | $9.5605e-10$ |
| | | | W-REM | $9.5605e-10$ |
|  |  |  | NREM-REM | 0.55869 |
| 128 Hz | 175.79 | 2, 166 | W-NREM | $9.5605e-10$ |
| | | | W-REM | $9.5605e-10$ |
| | | | NREM-REM | $9.5605e-10$ |
| 64 Hz | 234.46 | 2, 166 | W-NREM | $9.5605e-10$ |
| | | | W-REM | $9.5605e-10$ |
| | | | NREM-REM | $9.5605e-10$ |
| 32 Hz | 183.89 | 2, 166 | W-NREM | $9.5605e-10$ |
| | | | W-REM | $1.3432e-05$ |
| | | | NREM-REM | $9.561e-10$ |

**Table S.2.** Statistical comparisons between Amplitude Entropy values during sleep and wakefulness. This table provides the statistics for Fig. 5.

| Electrode | F | DF | W-NREM<br>(pValue) | W-REM<br>(pValue) | NREM-REM<br>(pValue) |
| --- | --- | --- | --- | --- | --- |
| OBr | 7.597 | 2,11 | 0.99 | 0.99 | 0.99 |
| M1r | 15.85 | 2,11 | 0.0001 | 0.70 | 0.03 |
| M1l | 12.49 | 2,11 | 0.0017 | 0.99 | 0.02 |
| S1r | 16.93 | 2,11 | 0.0016 | 0.06 | 0.09 |
| S1l | 14.63 | 2,11 | 0.0007 | 0.24 | 0.01 |
| V2r | 62.53 | 2,11 | 0.0003 | 0.0081 | 0.99 |
| V2l | 47.68 | 2,11 | 0.0004 | 0.0053 | 0.312 |

